# A non-canonical unfolded protein response pathway and mitochondrial dynamics control the number of ER-mitochondria contact sites

**DOI:** 10.1101/684753

**Authors:** Rieko Kojima, Yuriko Kakimoto, Manatsu Shinmyo, Kazuo Kurokawa, Akihiko Nakano, Toshiya Endo, Yasushi Tamura

**Affiliations:** Faculty of Science, Yamagata University, 1-4-12 Kojirakawa-machi, Yamagata 990-8560, Japan; Toyama Prefectural Institute for Pharmaceutical Research, 17-1 Nakataikouyama, Imizu, Toyama 939-0363, Japan; Department of Biochemistry and Molecular Biology, Graduate School of Medical Science, Yamagata University, 2-2-2 Iidanishi, Yamagata, 990-9585, Japan; Live Cell Super-Resolution Imaging Research Team, RIKEN Center for Advanced Photonics, 2-1 Hirosawa, Wako, Saitama 351-0198, Japan; Faculty of Life Sciences, Kyoto Sangyo University, Kamigamo-motoyama, Kita-ku, Kyoto 603-8555, Japan; Institute for Protein Dynamics, Kyoto Sangyo University, Kamigamo-motoyama, Kita-ku, Kyoto 603-8555, Japan

## Abstract

Mitochondria maintain their morphology and functions through the optimized balance between the mitochondrial fusion and division. Here we report a novel role of mitochondrial dynamics in controlling the number of ER-mitochondria encounter structure (ERMES) clusters in a yeast cell. Loss of mitochondrial fusion or division caused the increased or decreased number, respectively, of ERMES foci observed in cells. ERMES complexes, therefore, appear to cluster with each other and mitochondrial division may inhibit undesired ERMES hyper-clustering. Furthermore, our microscopic analyses suggest that ER stress induces dissociation of ERMES clusters, increasing the number of ERMES foci even in the absence of Ire1 and Hac1, which are essential factors for the UPR response. Interestingly, we found that ER stress leads to expansion of both the ER and mitochondrial membranes in an ERMES function-dependent manner. These findings imply that a cell is equipped with two independent regulatory mechanisms controlling the number of ER-mitochondria contact sites to meet the cellular as well as environmental demands.

## Introduction

Mitochondria are highly dynamic organelles and continuously fuse and divide to maintain their functionally optimized morphology. In yeast, two dynamin-related GTPases, Fzo1 and Mgm1 are known to mediate the mitochondrial outer and inner membrane (MOM and MIM) fusion, respectively [1]. Fzo1 is a MOM protein with two transmembrane (TM) segments and a large GTPase domain exposed to the cytosol. Mgm1 is present as two isoforms, long and short forms (l- and s-Mgm1). l-Mgm1 is integrated into the MIM via its first N-terminal TM segment and exposes its GTPase domain to the intermembrane space (IMS). s-Mgm1 is a soluble IMS protein, which is released from the MIM after cleavage of the long form by a rhomboid protease Pcp1 located in the MIM [2–5]. Another multi-spanning MOM protein Ugo1 physically interacts with both Fzo1 in the MOM and Mgm1 in the MIM and likely couples the MOM and MIM fusion events [6, 7]. Mitochondrial fusion is considered to contribute to attenuating mitochondrial oxidative damages, which could accumulate in mitochondria due to the generation of reactive oxygen species (ROS) as a byproduct of respiration. Impairment of the mitochondrial fusion leads to not only mitochondrial fragmentation by continuous mitochondrial fission but also to the loss of mitochondrial DNA (mtDNA). Consistent with the physiological importance of the mitochondrial fusion, mutations in Mfn2 and Opa1, mammalian Fzo1 and Mgm1, respectively, were found to cause human diseases such as Charcot-Marie-Tooth disease type 2A (CMT2A) and autosomal dominant optic atrophy (ADOA) [8]. Importantly, strong phenotypes associated with the cells defective in the mitochondrial fusion, growth defects as well as the loss of mtDNA can be suppressed by additional loss of division factors such as Dnm1 and Fis1 in yeast. Fragmented mitochondria in the cells with defective mitochondrial fusion are also restored to the ones in a tubular shape when mitochondrial division is additionally blocked [9–11].

Dnm1, another dynamin-related GTPase, is recruited to the MOM with the aid of receptor proteins such as Fis1, Mdv1, and Caf4 and assembled into a helical structure to split a mitochondrial tubule through its GTP hydrolysis-dependent conformational change [12–18]. Mitochondrial division in mammals further requires a classical dynamin, Dyn2, in addition to the mammalian counterpart of Dnm1, Drp1 (Lee et al., 2016). Mitochondrial division reportedly occurs at the mitochondria-ER contact sites, suggesting that the ERMES (ER-mitochondria encounter structure) complex, which directly tethers the ER to the MOM, may participate in mitochondrial division [19]. Mitochondrial tubules become constricted by physical association with the ER via the ERMES complex, and Dnm1 is targeted to such characteristic spots on mitochondria to mediate mitochondrial division [20].

The ERMES consists of four core subunits, ER-resident Mmm1, MOM-resident Mdm10 and Mdm34, and a peripheral membrane protein Mdm12 (Lang et al., 2015; Eisenberg-Bord et al., 2016; Murley and Nunnari, 2016; Tamura and Endo, 2017). These ERMES core subunits were initially identified as factors required for normal mitochondrial distribution and morphology [25–28]. Mitochondrial morphology is altered from a tubular to a spherical ball-like structure when one of these ERMES core components is absent, pointing out the close relationship of ERMES with mitochondrial morphogenesis. In addition to the role of ERMES in maintaining mitochondrial morphology, recent studies further revealed a function of ERMES in phospholipid transport between the ER and mitochondria (Kornmann et al., 2009; Kojima et al., 2016; Jeong et al., 2016, 2017; Kawano et al., 2018). These findings indicate that the ER-mitochondria tethering mediated by ERMES is vital for the functional integrity of mitochondria. Importantly, the ERMES complex constitutes large aggregates that can be observed as several discrete foci in a cell under a fluorescent microscope when an ERMES subunit is expressed as a GFP-fusion protein [25–28]. The number of ERMES in a cell should be precisely controlled to regulate the degree of mitochondrial division and phospholipid transfer. For example, the number of ERMES dots increases with a change in the metabolic state, such as a shift from the fermentation to non-fermentation culturing condition, which promotes mitochondrial proliferation [34]. Interestingly, physical association of mitochondria with the vacuole, termed vCLAMP (*v*a*c*uo*l*e *a*nd *m*itochondria *p*atch), exhibits an opposite behavior to ERMES with regard to the degree of membrane contacts; under non-fermentable conditions, Vps39, which is responsible for the formation of vCLAMP, is phosphorylated, resulting in a decreased number of vCLAMP [34]. These observations suggest that mitochondria-ER contacts and mitochondria-vacuole contacts mediated by ERMES and Vps39, respectively, are reciprocally regulated, and that the functions of ERMES and vCLAMP are partly overlapping [34, 35]. A high-content imaging screen for yeast deletion mutants with an altered number of ERMES dots revealed that loss of Vps39 or that of mitochondrial division factors such as Dnm1 and Fis1 leads to an increased number of ERMES dots [35]. These previous observations suggest that spatially separated inter-organelle contacts are functionally connected, yet their underlying mechanism remains to be elucidated.

Here, we analyzed the relationship of mitochondrial fusion and division with the number of ERMES dots and found that loss of mitochondrial fusion or division caused increased or decreased number, respectively. Besides, we found that treatment of cells with tunicamycin or DTT, which induces ER stress, led to dissociation of the ERMES clusters, increasing the number of ERMES dots, independently of the mitochondrial division as well as unfolded protein response (UPR) components, Ire1 and Hac1. More importantly, we found that both the ER and mitochondrial membranes enlarge upon ER stress in a Mmm1-dependent manner probably for relieving the ER stress. These findings strongly suggest that dynamic changes in the number and size of ER-mitochondria contact sites play an important role in dealing with the stress conditions.

## Results and Discussion

### Mitochondrial fusion and division antagonistically affect the number of ERMES foci

The main question we asked here is whether the cell has an active mechanism to regulate the number of inter-organelle contacts such as mitochondria-ER contact sites (MERCs), which are observed as dot-like structures under a microscope. If the cell is equipped with such a mechanism, we reasoned that regulators controlling the number of the MERCs could be localized in the MOM. To test this idea, we visualized MERCs with the split-GFP probes (Kakimoto et al., 2018) in yeast cells that lack each of the 53 different MOM proteins (Table S1). We noted that the number of MERCs significantly decreased in cells lacking Fis1, which functions as a receptor for mitochondrial division factor Dnm1 (Fig. S1) [14].

Since this observation is not consistent with the previous finding that the number of ERMES foci increased in the absence of Dnm1 or Fis1 [35], we further asked if the loss of mitochondrial division would affect the appearance of ERMES. We thus observed wild-type, *dnm1*Δ and *fis1*Δ cells expressing mitochondria-targeted RFP (Su9-RFP) and C-terminally GFP-tagged Mmm1 (Mmm1-GFP) under a confocal fluorescence microscope and acquired approximately 5-μm z-stack images that cover a whole cell with a 0.2-μm increment (Fig. 1). To minimize the undesired effects on the appearance of ERMES dots, arising from possible variations of the Mmm1-GFP expression level, we adopted the stable expression of Mmm1-GFP from the chromosome. Maximum projection images reconstituted from the z-stacks showed that ERMES dots were localized on mitochondrial tubules in wild-type, *dnm1*Δ, and *fis1*Δ cells although mitochondrial distribution was altered due to lack of mitochondrial division in *dnm1*Δ and *fis1*Δ cells; wild-type, *dnm1*Δ, and *fis1*Δ cells contained 5.8, 3.5 and 3.0 ERMES dots per cell on average, respectively. These results indicate that loss of mitochondrial division leads to a decrease in the number of ERMES dots. The reduced ERMES number observed in mitochondrial division-deficient cells was not due to decreased levels of ERMES subunits since the protein levels of Mmm1, Mdm12 and Mdm34 were all comparable between *dnm1*Δ cells and wild-type cells (Fig. S2A).

**Figure 1.**
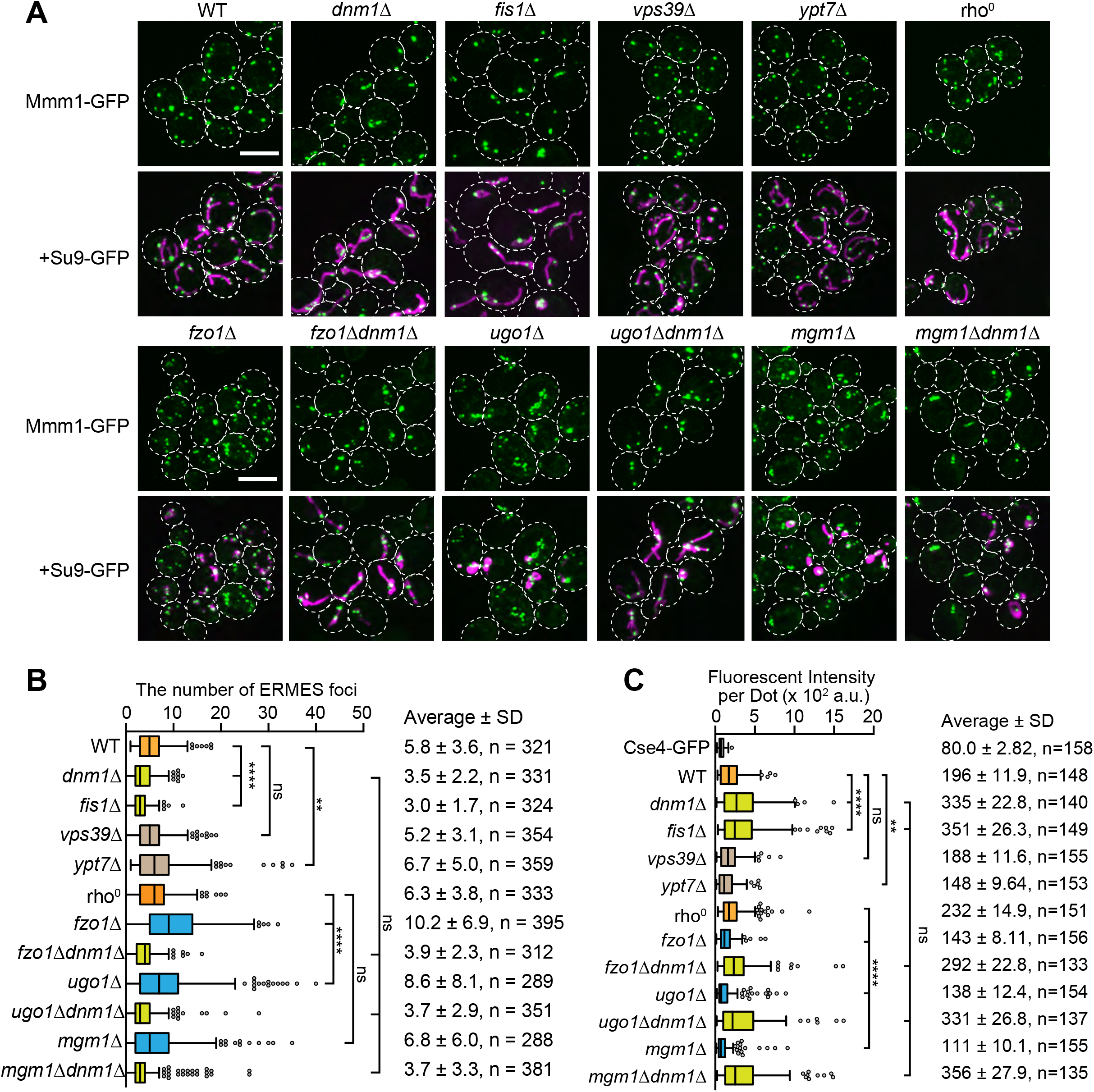
Mitochondrial fusion and division antagonistically regulate the number of ERMES dots. A The indicated yeast cells expressing Mmm1-GFP and mitochondria-targeted RFP (Su9-RFP) were imaged under a confocal fluorescence microscope. Maximum projection images were shown. Scale bars, 5 μm. B Box and whisker plots show the distribution of the number of ERMES dots per a cell. SD means standard deviation. **: *p* < 0.001,****: *p* < 0.0001, *p* values were obtained from the unpaired two-tailed t-test with Welch’s correction. C Quantifications of signal intensities of ERMES dots in the indicated cells using Cse4-GFP signal as a standard. Box and whisker plots show the estimated number of Mmm1-GFP molecules included in a single ERMES dot. The average intensity of Cse4-GFP foci was set to 80 U. ns: not significant, **: *p* < 0.001,****: *p* < 0.0001, *p* values were obtained from the unpaired two-tailed t-test with Welch’s correction.

We then re-examined the previously reported effects of loss of vCLAMP components on the number of ERMES foci. Although the vCLAMP component Vps39 was found as a factor whose absence led to an increase in the number of ERMES foci [35], we did not observe a drastic change in the number of ERMES foci in *vps39*Δ cells as compared with that of wild-type cells (Fig. 1A, B). On the other hand, loss of another vCLAMP component Ypt7 led to a slight increase in the ERMES dot number. These results collectively suggest that factors other than vCLAMP, e.g., the growth phase, growth media, or yeast strain backgrounds could also affect the appearance of ERMES dots in a complex manner.

We next estimated the number of Mmm1-GFP molecules included in a single ERMES dot from their GFP signals using Cse4-GFP as a standard [36, 37]. Interestingly, we found that each of the ERMES dots in *dnm1*Δ and *fis1*Δ cells contains 335 and 351 of Mmm1-GFP molecules on average, respectively, which are significantly larger than 196 molecules per dot in wild-type cells (Fig. 1C). The average numbers of Mmm1-GFP molecules per ERMES dot were similar in *vps39*Δ and slightly smaller in *ypt7*Δ cells as compared with that in wild-type cells. These results suggest that the reduced number of ERMES dots in yeast mutant cells defective in mitochondrial division reflects enhanced clustering of the preexisting ERMES clusters. The proper mitochondrial division may thus inhibit unnecessary clustering of ERMES complexes, thereby maintaining the appropriate number of the MERCs in cells.

If mitochondrial division suppresses hyper-clustering of preexisting ERMES clusters and thereby controls the optimum number of ERMES foci, loss of mitochondrial fusion will, in turn, increase the number of ERMES foci due to ongoing mitochondrial division. We thus deleted mitochondrial fusion genes, *FZO1*, *UGO1* or *MGM1* and observed ERMES dots visualized with Mmm1-GFP. Since the lack of mitochondrial fusion causes loss of mtDNA, we used a *rho*^0^ strain, in which mitochondrial DNA lacks, as a control. Supporting our above idea, defects in mitochondrial fusion by deletion of the genes for the MOM fusion, *FZO1*, and *UGO1*, increased the number of ERMES foci while the loss of mtDNA alone did not (Fig. 1A, B). Deletion of the MIM fusion gene, *MGM1*, caused a marginal increase in the ERMES dot number, which could reflect the fact that Mgm1-mediated MIM fusion is not entirely coupled with the Fzo1-mediated MOM fusion [38]. Further supporting the role of mitochondrial division in controlling the number of ERMES foci, loss of the division gene *dnm1*Δ together with the loss of a fusion gene, *FZO1*, *UGO1*, or *MGM1* reversed the phenotypes. That is, *fzo1*Δ*dnm1*Δ, *ugo1*Δ*dnm1*Δ, and *mgm1*Δ*dnm1*Δ cells exhibited tubular mitochondria due to simultaneous defects in mitochondrial fusion and division as reported, and the number of ERMES foci was changed to the level similar to the one for *dnm1*Δ cells, regardless of the simultaneous deletion of *FZO1*, *UGO1* or *MGM1* (Fig. 1A, B). Besides, the estimated number of Mmm1-GFP molecules in an ERMES dot was clearly smaller when mitochondrial fusion was inhibited than those of wild-type and *rho*^*0*^ cells while the Mmm1-GFP particles per ERMES dot drastically increased to the level similar to *dnm1*Δ cells when *dnm1*Δ was deleted together with *FZO1*, *UGO1* or *MGM1* (Fig. 1C). These results suggest that mitochondrial fusion and division antagonistically affect the number of ERMES foci by controlling the clustering of preexisting ERMES foci.

### Preexisting ERMES foci cluster together

To directly test the idea that the preexisting ERMES foci gather together, we utilized a yeast mating assay with which we can monitor mitochondrial fusion directly (Sesaki et al., 2003b) (Fig. 2A). Briefly, we constructed two types of yeast cells with the opposing mating types (*MAT*a and *MAT*α) that express Mmm1-GFP or Mmm1-mScarlet under the control of the *GAL1* promoter. Then we examined how the ERMES foci behave after cell fusion. If the preexisting ERMES foci tend to cluster together, the green- and red-labeled foci should be merged or adjacent to each other after cell fusion (Fig. 2A). To exclude the mere possibility that newly synthesized Mmm1-GFP congregates into preexisting ERMES foci consisting of Mmm1-mScarlet, or vice versa, after cell fusion, we suppressed the expression of Mmm1-GFP and Mmm1-mScarlet before mating by cultivating the cells in a glucose-containing medium YPD for 4 hours. Immunoblotting of whole cell lysates confirmed that Mmm1-GFP and Mmm1-mScarlet expression was shut off after cultivation in YPD (Fig. 2B). Strikingly, most ERMES foci labeled with different fluorescent proteins were adjacent to each other rather than completely merged (Fig. 2C). This observation indicates that the different preexisting ERMES foci do not completely fuse, but instead contact with each other to form large clusters. We thus propose that the proper balance of mitochondrial fusion and division regulates clustering of preexisting ERMES foci, thereby maintaining the optimum number of ERMES foci (Fig. 2D).

**Figure 2.**
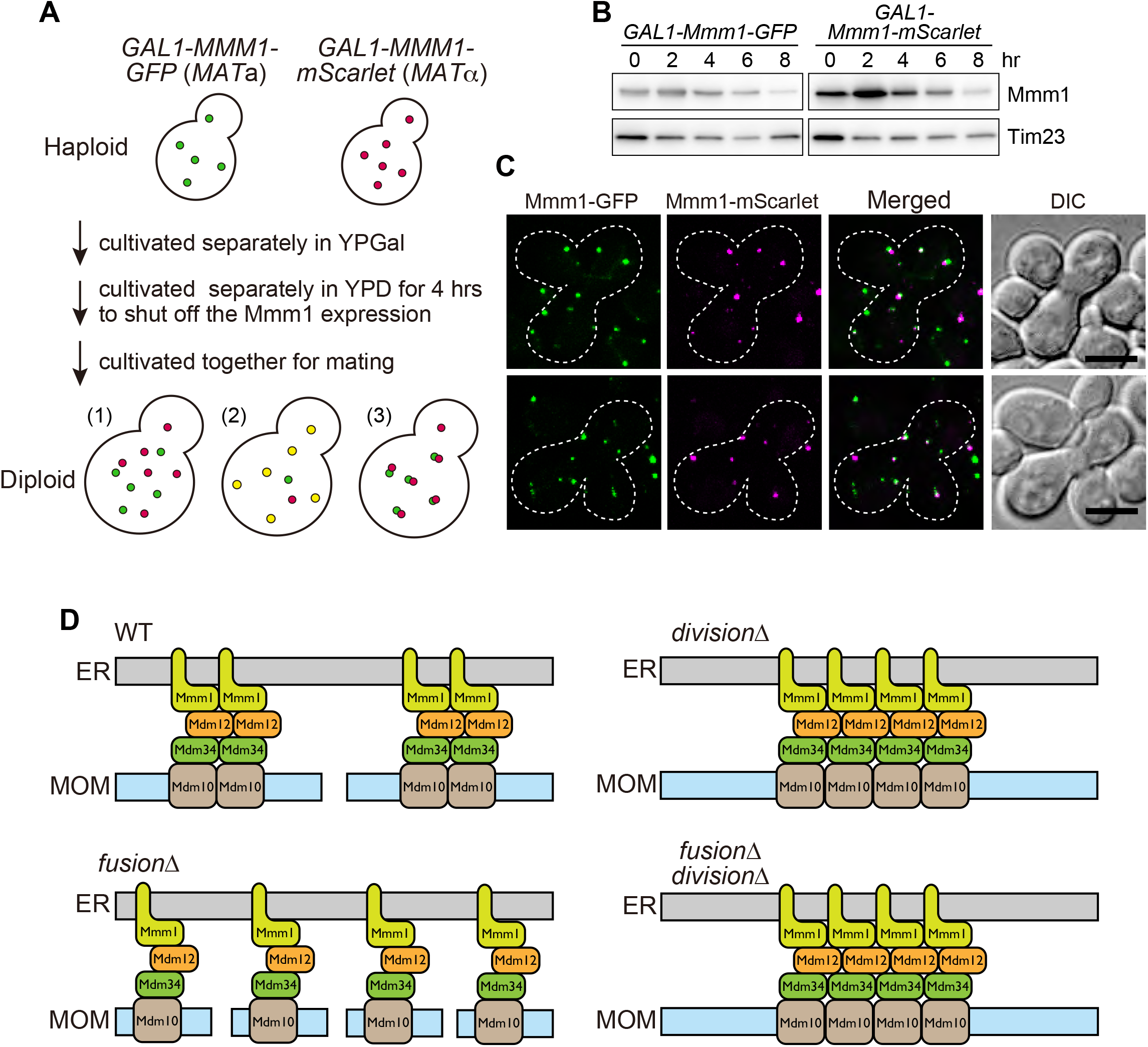
Pre-existing ERMES dots cluster together upon cell fusion. A A schematic diagram of the mating assay showing possible patterns of the ERMES dots. (1) The preexisting ERMES foci do not cluster together. (2, 3) The preexisting ERMES foci merge completely (2) or contact each other (3). B Whole cell extracts prepared from yeast cells after shutting off the expression of Mmm1-GFP or Mmm1-mScarlet were subjected to immunoblotting using antibodies against Mmm1 and Tim23. C Yeast zygotes obtained by mating haploid cells containing ERMES dots labeled with different fluorescent colors. Scale bars, 5 μm. D A working model for regulating the number of ERMES dots by mitochondrial fusion and division.

### The ERMES foci number increases upon ER stress in an Ire1- and Hac1-independent manner

Studies using mammalian cultured cells suggested that the MERC number increased under ER stress conditions [39–41]. In contrast, a recent study reported that ER stress did not drastically affect MERC foci visualized with split-GFP probes in U2OS cells [42]. We thus treated yeast cells expressing Mmm1-GFP with tunicamycin or DTT for 2 hours to induce ER stress, and then observed ERMES foci under a fluorescence microscope. Strikingly, the number of ERMES foci increased about 2-fold upon treatment with tunicamycin or DTT (Fig. 3A, C). We confirmed that tunicamycin treatment does not affect the amounts of ERMES components such as Mmm1 and Mdm12 although it decreased the level of N-glycosylated Mmm1 and Mmm1-GFP (Fig. S2B). Therefore, the sharp increase in the number of ERMES foci was not due to increased expression of ERMES components. The growth in the ERMES dot number upon ER stress was not due to accelerated mitochondrial division, either, since tunicamycin or DTT treatment caused an increase in the number of ERMES foci even in the absence of Dnm1 (Fig. 3B, C). Another possible explanation for the increased number of ERMES foci is that the ERMES clusters dissociate upon ER stress. We examined this possibility by super-resolution confocal live imaging microscopy (SCLIM), which enable us to obtain time-lapse fluorescent images of whole yeast cells at super-high resolution [43, 44]. Interestingly, we could observe splits of the ERMES dots after the tunicamycin treatment (Fig. S2C). Consistently, we confirmed that the estimated number of Mmm1-GFP molecules present in an ERMES dot after the tunicamycin or DTT treatment is smaller than that of nontreated wild-type cells (Fig. 3D). These findings strongly suggest that dissociation of ERMES clusters is at least partly responsible for the increased ERMES foci number.

**Figure 3.**
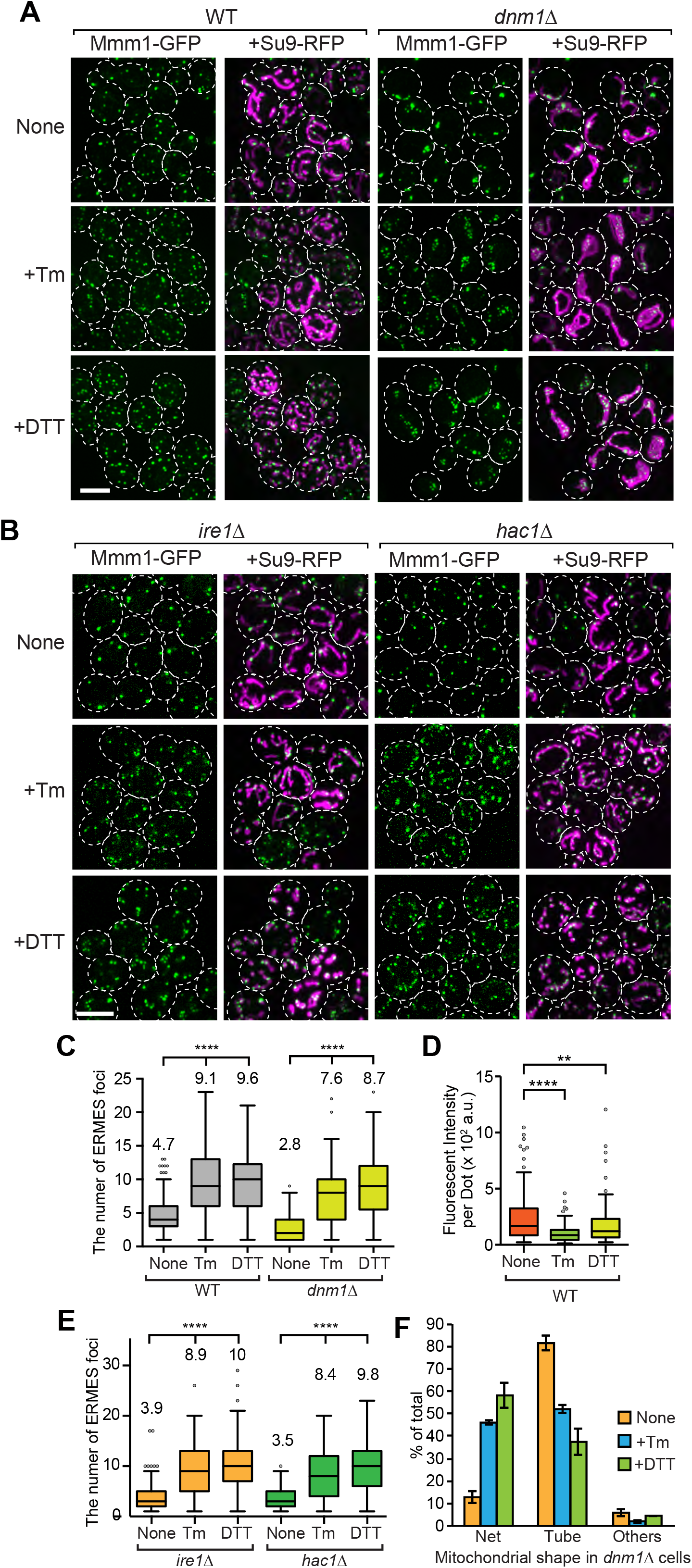
The number of ERMES dots significantly increases upon treatments with ER stress inducers independently of mitochondrial division and the conventional UPR pathway. A Wild-type (WT) and *dnm1*Δ cells expressing Mmm1-GFP and mitochondria-targeted RFP (Su9-RFP) were incubated in SCD media with or without 1 μg/ml tunicamycin (+Tm) or 3 mM DTT (+DTT) for 2 h and then observed under a confocal fluorescence microscope. Maximum projection images reconstituted from z-stacks were shown. Scale bars, 5 μm. B *ire1*Δ and *hac1*Δ cells expressing Mmm1-GFP and mitochondria-targeted RFP (Su9-RFP) were observed under a confocal fluorescence microscope as in A. C Box and whisker plots show the number of ERMES dots per a wild-type or a *dnm1*Δ cell with or without the treatment inducing ER stress. Numbers indicated above the plots show mean. n = 199, 194 and 132 (WT, None, Tm and DTT) and n = 179, 174 and 105 (*dnm1*Δ, None, Tm and DTT). ****: *p* < 0.0001, *p* values were obtained from the unpaired two-tailed t-test with Welch’s correction. D Quantifications of signal intensities of ERMES dots in wild-type cells with or without the DTT or Tm treatment as in Fig. 1C. **: *p* = 0.0019, ****: *p* < 0.0001, *p* values were obtained from the unpaired two-tailed t-test with Welch’s correction. E The number of ERMES dots per an *ire1*Δ or a *hac1*Δ cell with or without the treatment inducing ER stress was quantified as in Fig. 4C. n = 386, 252 and 315 (*ire1*Δ, None, +Tm and +DTT) and n = 386, 323 and 300 (*hac1*Δ, None, +Tm and +DTT). ****: *p* < 0.0001, *p* values were obtained from the unpaired two-tailed t-test with Welch’s correction. F Quantifications of the ratio of cells containing large net-like (Net), tubular (T), and the other shaped mitochondria in Fig. 3B. Total 436, 451 and 245 cells for None (without inducing ER-stress), +Tm (with tunicamycin treatment) or +DTT (with DTT treatment), respectively, were examined in three independent experiments. Values are mean±S.E.(n = 3)

We next asked whether the increase in the number of ERMES foci would depend on the unfolded protein response (UPR) [45]. In yeast, an ER-resident type I TM protein Ire1 senses accumulation of aberrant proteins in the ER. Then Ire1 undergoes self-oligomerization, which triggers both kinase and endonuclease activation of Ire1, hence leading to Ire1-mediated *HAC1* mRNA splicing [46]. The spliced mature *HAC1* mRNAs produce the functional form of the Hac1 transcription factor, which activates a number of genes repressed under non-stress conditions [47, 48]. We thus asked if a loss of either Ire1 or Hac1 would affect the increase in the number of ERMES foci under ER stress conditions. Surprisingly, neither loss of Ire1 nor Hac1 suppressed the increase in the number of ERMES foci (Fig. 3B, F), suggesting that a non-canonical mechanism independent of the *IRE1*/*HAC* mediates the ER-stress dependent change in the ERMES foci number. What mechanism underlies the dissociation of ERMES clusters under ER stress conditions? One possibility is that ER stress leads to dissociation of ERMES clusters through modifications like phosphorylation and ubiquitination of the ERMES components. However, this is unlikely because we did not observe band shifts of any of the ERMES components by SDS-PAGE followed by immunoblotting (Fig. S2C). We also confirmed that yeast cells expressing Mdm34-3PA mutant, which lacks the PY motif critical for its ubiquitination by Rsp5 E3 ubiquitin ligase, showed a similar increase in the ERMES dot number after tunicamycin and DTT treatments (Fig. S2D). Another possibility is that the ER stress induces alteration of phospholipid compositions of the ER membrane and/or MOM, which may affect the conformation and/or assembly of the ERMES complex, resulting in the split of the ERMES foci. However, we confirmed that at least phospholipid class compositions were not altered by tunicamycin and DTT treatments (data not shown).

### ER-stress leads to expansion of both the ER and mitochondria membranes through ERMES functions

What is the possible physiological role of the ER stress-triggered increase in the ERMES dot number? A previous study showed that the ER membranes significantly expand upon ER stress in an Ino2/Ino4-dependent manner to attenuate the ER stress [49]. Ino2/Ino4 transcription factors are known to activate transcriptions of a series of genes for phospholipid synthases [50]. On the other hand, it is well known that phospholipids have to shuttle between the ER and mitochondria via ERMES for their proper syntheses [51–54]. Therefore, it is attractive to assume that the combination of transcriptional activation of the genes for phospholipid synthetic enzymes and the increase in the number of ERMES dots, which represent phospholipid transport sites between the ER and mitochondria, cooperatively enhance phospholipid biosynthesis under ER stress conditions. Supporting this idea, we found that the ER stress-dependent ER membrane expansion is partly suppressed by loss of Mmm1 (Fig. 4A). ~90% of wild-type cells showed the ER structure with elongated membranes after ER stress inductions whereas ~50% of *mmm1*Δ cells contained such a developed ER structure (Fig. 4C). More importantly, we found that tunicamycin and DTT treatments led to a significant increase in mitochondrial structure as well (Fig. 4B, D). We noticed that the loss of Mmm1 abolished the development of mitochondria membranes under these ER stress conditions. Similar mitochondrial expansion was observed when mitochondrial division factor Dnm1 is absent. It is known that the lack of a division factor Dnm1 leads to a change in mitochondrial morphology from tubular structures to net-like structures [9, 10]. The net-like mitochondrial shape was further enhanced likely owing to the mitochondrial membrane expansion by the ER stress (Fig. 3A, F). There results suggest that expansion of the ER and mitochondria membranes requires normal ERMES functions. Consistent with this idea, we found that *mmm1-1* cells became more susceptible to the tunicamycin treatment as compared with the corresponding wild-type strain (Fig, 4E). Collectively, our results strongly suggest a previously overlooked fact that ER stress induces the Ire1/Hac1-independent dissociation of ERMES clusters, which could activate phospholipid biogenesis in cooperation with the Ire1/Hac1-dependent transcriptional upregulation of phospholipid synthase genes. The enhanced phospholipid biogenesis would be critical for expansion of not only the ER but also mitochondrial membranes to buffer the ER stress (Fig. 4F).

**Figure 4.**
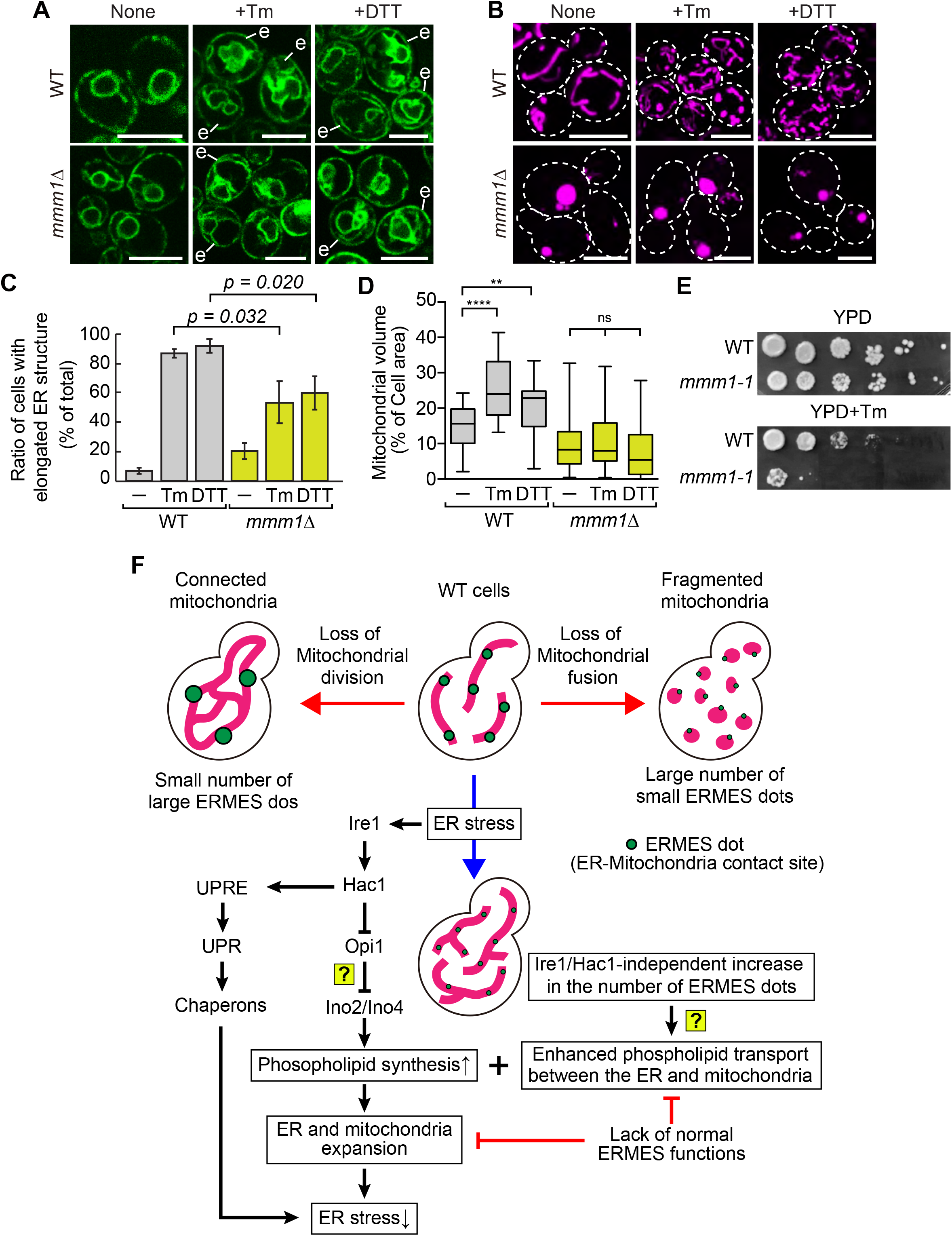
ER stress causes membrane expansion of both the ER and mitochondria in a Mmm1-dependent manner. A Wild-type and *mmm1*Δ cells expressing the ER-targeted or GFP were imaged under a confocal fluorescence microscope. Maximum projection images were shown. Scale bars, 5 μm. “e” in (A) represents the expanded ER structure. B Wild-type and *mmm1*Δ cells expressing the mitochondria-targeted GFP were imaged as in A. C Ratio of cells containing the expanded ER membranes. A Total of over 180 cells were evaluated by three independent experiments. Values are mean±S.E. (n = 3) D Box and whisker plots show relative area of mitochondrial membranes to cell area. **: *p* = 0.0019, ****: *p* < 0.0001, *p* values were obtained from the unpaired two-tailed t-test with Welch’s correction. E *mmm1-1* and its corresponding wild-type cells were spotted onto YPD with or without Tm and cultivated at 23°C for 6 days. F A schematic showing a model that combination of canonical and non-canonical UPR response pathways cooperatively contribute to membrane expansion of the ER and mitochondria to cope with ER stress.

However, there are some unresolved issues here. For example, it is still unclear how Ino2/Ino4 transcription factors are activated in Ire1/Hac1-dependent manner upon ER stress. A previous study suggested a possibility that Hac1 inhibits the function of Opi1 as a repressor for Ino2/Ino4 although this has not yet been revealed (Fig. 4F). Whether the lipid flux between the ER and mitochondria is really activated under ER stress conditions is also an important question to be assessed. Besides, what triggers the dissociation of ERMES clusters remains unclear. Although we confirmed that ER stress induction does not alter compositions of at least head group of phospholipids, it is still possible that acyl-chain variations cause conformational changes in the ERMES clusters, which could lead to their dissociation. Alternatively, protein modifications or additional binding partner for ERMES subunits may cause the dissociation of ERMES clusters under ER stress conditions. Detail mechanisms underlying the dynamic feature of the number of ERMES clusters should be investigated in future studies

## Material and Methods

### Strains, plasmids, primers, and growth conditions

In this study, we used a *Saccharomyces cerevisiae* strain, FY833 (MATa *ura3-52 his3*-Δ*200 leu2*-Δ*1 lys2*-Δ*202 trp1*-Δ*63*) or FY834 (MATα *ura3-52 his3*-Δ*200 leu2*-Δ*1 lys2*-Δ*202 trp1*-Δ*63*) as background strains [55]. All the yeast cells used in this study are listed in Table S1. Yeast cells expressing Mmm1-GFP originated from FY833 was used as a wild-type strain throughout this study [55, 56]. The C-terminal GFP or mScarlet tagging, the introduction of the *GAL1* promoter in front of the *MMM1* gene and gene disruptions were performed by homologous recombination using the appropriate gene cassettes amplified from plasmids listed in Table S2 [57, 58]. A pair of primers, #YU1116/1117, #NU831/832, #NU829/830, #YU29/30, #YU31/32, #YU33/34, #YU1347/1348, or #YU1349/1350 was used to amplify the gene cassette for the disruption for *FIS1*, *VPS39*, *YPT7*, *FZO1*, *UGO1*, *MGM1*, *IRE1* or *HAC1* gene, respectively. For the introduction of GFP or Scarlet tag for *MMM1 and CSE4*, the *GAL1* promoter for *MMM1* or 3PA-GFP tag for *MDM34*, the appropriate gene cassettes were amplified with a pair of primer #NU1174/1175, #YU1518/1519, #YU1563/1564, or #NU1071/1072, respectively. Yeast cells were grown in YPD (1% yeast extract, 2% polypeptone, and 2% glucose), SCD (0.67% yeast nitrogen base without amino acids, 0.5% casamino acid, and 2% glucose), SD (0.67% yeast nitrogen base without amino acids, 0.13% drop-out amino acid mix and 2% glucose) media with appropriate supplements. The drop-out amino acid mix was a mixture of 2.6 g adenine, 6.0 g L-aspartic acid, 12 g L-threonine, 2.6 g L-asparagine, 1.8 g L-tyrosine, 6.0 g L-glutamic acid, 2.6 g L-glutamine, 2.6 g glycine, 2.6 g L-alanine, 2.6 g L-isoleucine, 1.2 g L-methionine, 3.0 g L-phenylalanine, 2.6 g L-proline, 22.6 g L-serine, 9.0 g L-valine and 2.6 g L-cysteine.

### Fluorescence microscopy

Logarithmically growing yeast cells cultivated in SCD or SD media were observed under Olympus IX83 microscope with a CSU-X1 confocal unit (Yokogawa), a 100 x, 1.4 NA, objective (UPlanSApo, Olympus) and an sCMOS camera (Zyla 5.5; Andor) manipulated by MetaMorph software (Molecular Devices). When observed mScarlet signal, we used an EM-CCD camera (Evolve 512; Photometrics). GFP or RFP/mScarlet were excited by 488-nm or 561-nm laser (OBIS, Coherent) and the emission was passed through 520/35-nm or 617/73-nm band-pass filter, respectively. The confocal fluorescent sections were collected every 0.2 μm from the upper to the bottom surface of yeast cells. The obtained confocal images were subjected to maximum projection using Image J software. For counting ERMES dots, Mmm1-GFP dots were automatically picked by TransFluor, a macro for MetaMorph software, using the “Pits” algorithm. The resulting images containing picked dots were then subjected for counting. Signal intensities of ERMES and Cse4-GFP dots were calculated by using Image J software.

SCLIM was developed by combining Olympus model IX-71 inverted fluorescence microscope with a UPlanSApo 100 X NA 1.4 oil objective lens (Olympus, Japan), a high-speed and high-signal-to noise-ratio spinning-disk confocal scanner (Yokogawa Electric, Japan), a custom-made spectroscopic unit, image intensifiers (Hamamatsu Photonics, Japan) equipped with a custom-made cooling system, magnification lens system for giving 266.7 X final magnification, and three EM-CCD cameras (Hamamatsu Photonics, Japan) for green, red, and infrared observation [43]. Image acquisition was executed by custom-made software (Yokogawa Electric, Japan). For 3D time-lapse imaging, we collected optical sections spaced 0.2 μm apart in stacks by oscillating the objective lens vertically with a custom-made piezo actuator. Z stack images were converted to 3D voxel data and processed by deconvolution with Volocity (Perkin Elmer, MA) using the theoretical point-spread function for spinning-disk confocal microscopy.

### Immunoblotting and antibodies

For immunoblotting, proteins transferred to PVDF membranes (Immobilon-FL or Immobilon-P, Millipore) were detected by fluorophore- or HRP-conjugated to secondary antibodies (Cy5 AffiniPure Goat Anti-Rabbit IgG (H+L) from Jackson ImmunoResearch Labs or Goat anti-Rabbit IgG (H+L) Cross-Adsorbed Secondary Antibody, HRP from Thermo Fisher Scientific) and analyzed with a Typhoon imager (GE Healthcare) or LAS-4000 mini (Fujifilm).

### Statistical analyses

The number of ERMES per a cell and signal intensities of ERMES and Cse4-GFP dots were shown as box and whisker plots (Tukey). The sample number was shown in figure legends. Data are shown as means with SEM or SD as indicated in the figure legends. The Student’s t-test with Welch’s correction were performed for the statistical analyses using Prism 6 (GraphPad).

## Acknowledgments

We thank K. Shishido, M. Hashimoto, and T. Sasaki for their great technical assistance and Profs. Robert E. Jensen and Hiromi Sesaki for *mmm1-1* and its corresponding wild-type strain. We are grateful to the members of the Tamura and Endo laboratories for helpful discussion. This work was supported by JSPS KAKENHI (Grant Numbers 17H06414 and 19H03174 to YT, 17H06413 to KK and AN, 17H06420 and 18H05275 to K.K, and 15H05705 and 22227003 to TE), AMED-PRIME (Grant Number JP19gm5910026) from Japan Agency for Medical Research and Development, AMED (YT), and a CREST Grant (JPMJCR12M1) from JST (TE).

## Author Contributions

TE, RK, and YT designed the study. RK performed the experiments shown in Fig. 1 and 2. YK performed the analysis shown in Fig. 4. SM screened for MOM proteins whose loss affected the appearance of the ER-mitochondria contact sites visualized with split-GFP probes. KK and AN helped to conduct live-cell imaging with SCLIM and edited the manuscript. TE and YT wrote the paper.

