## Supplemental information for "A non-canonical unfolded protein response pathway and mitochondrial dynamics control the number of ER-mitochondria contact sites"

**Figure S1.** Loss of Fis1 leads to decreased ER-mitochondria contact sites visualized by split-GFP probes. Related to Figure 1.

**Figure S2.** Steady state levels of ERMES components are not altered in cell lacking mitochondrial fusion and division factors. Related to Figure 2.

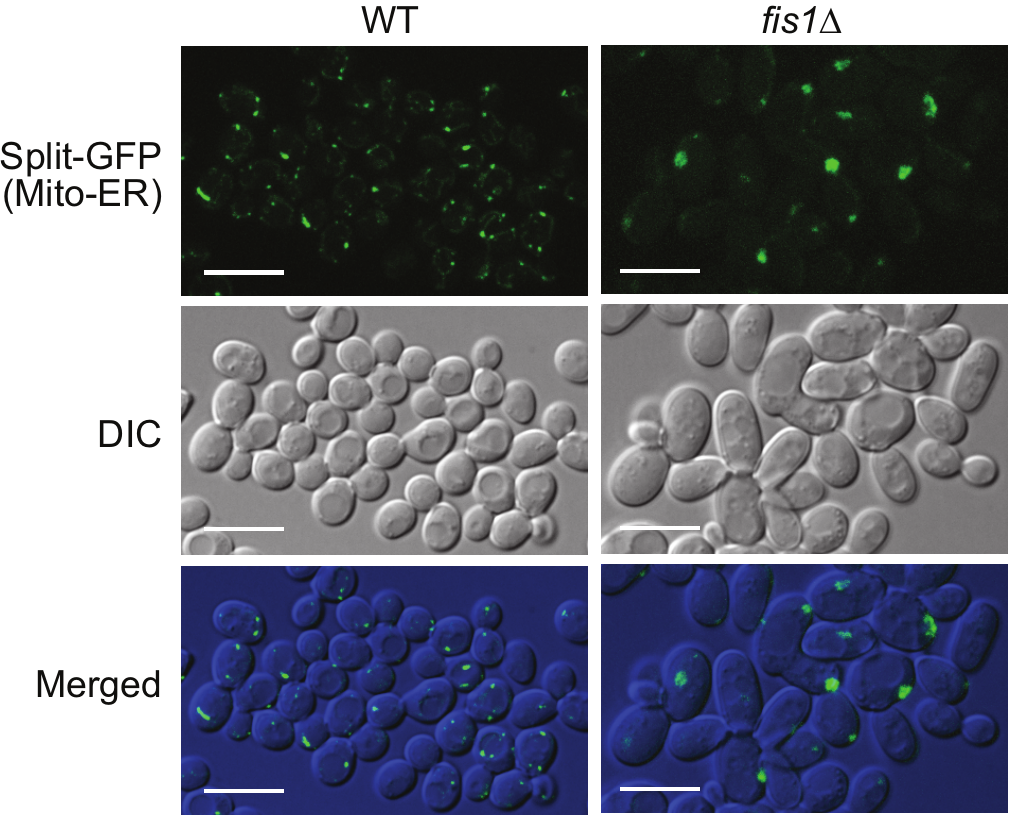

**Figure S1.** Loss of Fis1 leads to decreased ER-mitochondria contact sites visualized by split-GFP probes. Related to Figure 1.

Wild-type and *fis1*∆ cells expressing split-GFP probes (Tom71-V5-GFP11 and Ifa38-GFP1-10) were observed under a confocal fluorescence microscope. Maximum projection images were shown. Scale bars, 10 µm.

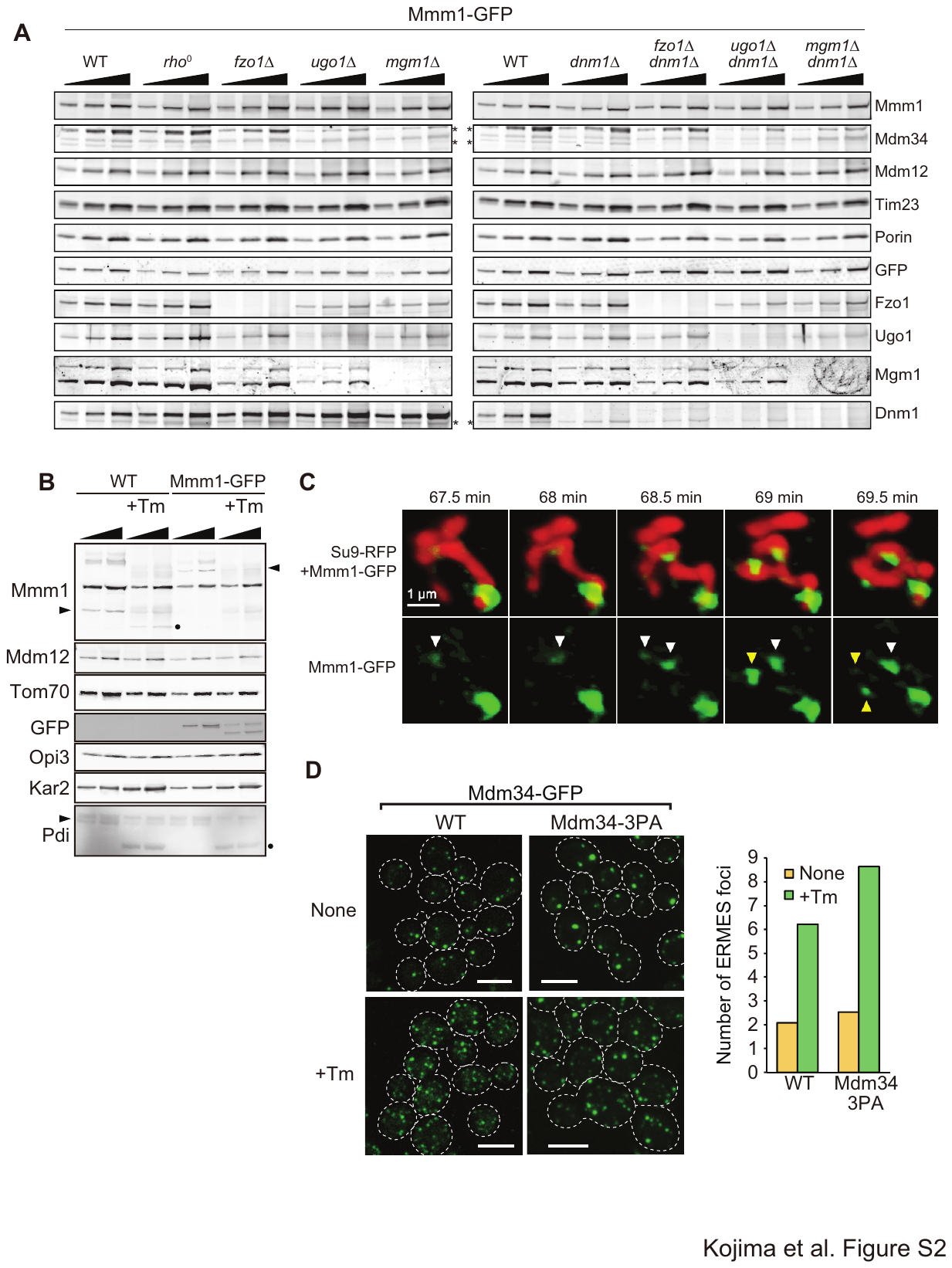

**Figure S2.** Steady state levels of ERMES components are not altered in cell lacking mitochondrial fusion and division factors. Related to Figure 2.

(A, B) Membrane fractions prepared from the indicated cells with or without tunicamycin or DTT treatment were analyzed by immunoblotting using the indicated antibodies. *, nonspecific bands. Arrow heads indicate Mmm1, Mmm1-GFP and Pdi bands. Black dot indicates non-glycosylated Mmm1 band. (C) Time-lapse images of whole yeast cells expressing Mmm1-GFP and Su9-RFP were taken by SCLIM microscope. Triangles shows ERMES dots that divide during the observation. Scale bar, 1 µm. (D) Yeast cells expressing Mdm34-GFP or Mdm34-3PA-GFP were imaged by confocal fluorescence microscope. Scale bars, 5 µm.
