## Supplementary material for "A non-canonical unfolded protein response pathway and mitochondrial dynamics control the number of ER-mitochondria contact sites": Table S1, S2

| Table S1 | |
| --- | --- |
| All listed below are *Saccharomyces cerevisiae* strains | Source |
| FY833- MATa *ura3-52 his3-∆200 leu2-∆1 lys2-∆202 trp1-∆63* | Winston et al., 1995 |
| FY834- MATα *ura3-52 his3-∆200 leu2-∆1 lys2-∆202 trp1-∆63* | Winston et al., 1995 |
| *Mmm1-GFP-* MATa *ura3-52 his3-∆200 leu2-∆1 lys2-∆202 trp1-∆63 MMM1-GFP::TRP1* | Tamura et al., 2012 |
| *Mmm1-GFP dnm1∆-* MATa *ura3-52 his3-∆200 leu2-∆1 lys2-∆202 trp1-∆63 MMM1-GFP::TRP1 dnm1∆::URA3* | This study |
| *Mmm1-GFP vps39∆-* MATa *ura3-52 his3-∆200 leu2-∆1 lys2-∆202 trp1-∆63 MMM1-GFP::TRP1 vps39∆::hphMX* | This study |
| *Mmm1-GFP ypt7∆-* MATa *ura3-52 his3-∆200 leu2-∆1 lys2-∆202 trp1-∆63 MMM1-GFP::TRP1 ypt7∆::hphMX* | This study |
| *Mmm1-GFP* ρ^o^- MATa *ura3-52 his3-∆200 leu2-∆1 lys2-∆202 trp1-∆63 MMM1-GFP::TRP1* ρ^o^ | This study |
| *Mmm1-GFP* *fzo1∆*- MATa *ura3-52 his3-∆200 leu2-∆1 lys2-∆202 trp1-∆63 MMM1-GFP::TRP1 fzo1∆::kanMX4* ρ^o^ | This study |
| *Mmm1-GFP* *fzo1∆dnm1∆*- MATa *ura3-52 his3-∆200 leu2-∆1 lys2-∆202 trp1-∆63 MMM1-GFP::TRP1 fzo1∆::kanMX4 dnm1∆:: kanMX4* | This study |
| *Mmm1-GFP* *ugo1∆*- MATa *ura3-52 his3-∆200 leu2-∆1 lys2-∆202 trp1-∆63 MMM1-GFP::TRP1 ugo1∆::HIS3* ρ^o^ | This study |
| *Mmm1-GFP* *ugo1∆dnm1∆*- MATa *ura3-52 his3-∆200 leu2-∆1 lys2-∆202 trp1-∆63 MMM1-GFP::TRP1 ugo1∆::HIS3 dnm1∆:: kanMX4* | This study |
| *Mmm1-GFP* *mgm1∆*- MATa *ura3-52 his3-∆200 leu2-∆1 lys2-∆202 trp1-∆63 MMM1-GFP::TRP1 mgm1∆::kanMX4* ρ^o^ | This study |
| *Mmm1-GFP* *mgm1∆dnm1∆*- MATa *ura3-52 his3-∆200 leu2-∆1 lys2-∆202 trp1-∆63 MMM1-GFP::TRP1 mgm1∆::kanMX4 dnm1∆:: kanMX4* | This study |
| *Mmm1-GFP* *ire1∆*- MATa *ura3-52 his3-∆200 leu2-∆1 lys2-∆202 trp1-∆63 MMM1-GFP::TRP1 ire1∆::kanMX4* | This study |
| *Mmm1-GFP* *hac1∆*- MATa *ura3-52 his3-∆200 leu2-∆1 lys2-∆202 trp1-∆63 MMM1-GFP::TRP1 hac1∆::kanMX4* | This study |
| *GAL-Mmm1-GFP*- MATa *ura3-52 his3-∆200 leu2-∆1 lys2-∆202 trp1-∆63 His3MX6-GAL1-MMM1-GFP::TRP1* | This study |
| *GAL-Mmm1-mScarlet*- MATα *ura3-52 his3-∆200 leu2-∆1 lys2-∆202 trp1-∆63 His3MX6-GAL1-MMM1-mScarlet::kanMX4* | This study |
| *Mmm1-3PA-GFP*- MATa *ura3-52 his3-∆200 leu2-∆1 lys2-∆202 trp1-∆63 MMM1-3PA-GFP::TRP1* | This study |
| *Cse4-GFP*- MATa *ura3-52 his3-∆200 leu2-∆1 lys2-∆202 trp1-∆63 Cse4-GFP::TRP1* | This study |
| YPH250*-MATa ura3-52 lys2-801 ade2-101 trp1-Δ1 his3-Δ200 leu2-Δ1 gal3* | Sikorski and Hieter, 1989 |
| *mmm1-1- MATa ura3-52 lys2-801 ade2-101 trp1-Δ1 his3-Δ200 leu2-Δ1 gal3 mmm1-1* | Burgess et al., 1994 |
| Table S2. |  |
| Sequence | Primer Name |
| AACATTATCTGATATCACGGATAGAGGCAAAACGGTAGGCTCATTTAACGGTTGTAAAACGACGGCCAGT | YU29 |
| TAACATTATGTATATTGATTTGAAAAGACCTCATATATTTACAAGAATATCACAGGAAACAGCTATGACC | YU30 |
| TCTCTGCCGATTTTTGGTTTCCAATAGTATAGGTTTAACTCAACCCCCCAGTTGTAAAACGACGGCCAGT | YU31 |
| CACTGGAATACCATGGGCGAACATAAAAAAAAATGGGGGACTATCCCAGTCACAGGAAACAGCTATGACC | YU32 |
| TTTCGTACTTATAATATAGCCTCGCATATTCACCATAATACTCTGAAAGCGTTGTAAAACGACGGCCAGT | YU33 |
| CGGTAAAAAATGCTATTTACAAATTCTCTAATGACACTATTTATTTTACACACAGGAAACAGCTATGACC | YU34 |
| GACGGGTCTTATATTGATCAGCAAAAACCCTTCAAAATATCAATTTATACCAAAAATTAAGTTGTAAAACGACGGCCAGT | NU831 |
| GAGAGATTTTTAATATATATAAGAAATACTAACAACAATAACAGCAGCTGTTAAGGGATCCACAGGAAACAGCTATGACC | NU832 |
| GTTACCAAGTATGTGGCCACGTAGTAAAAATACGAGAGAAGAAAAGCCTACAGAGTTACGGATCCCCGGGTTAATTAA | NU1174 |
| CCAAAAATGAGGCAGAGAAGATAGGAAAAAGATAGAACAAAAAATTTGTACATAAATATGAATTCGAGCTCGTTTAAAC | NU1175 |
| TGAAAGAACTTTGAGAGAGTCAATATAATACCTGTAGCCTTTTTCTGAAAGAATTCGAGCTCGTTTAAAC | YU1071 |
| CAAACGTCATTAACGAATCCGTTTCGGTGGATTCATTCTCACTATCAGTCATTTTGAGATCCGGGTTTT | YU1072 |
| CATAGAAGCACAGATCAGAGCACAGCCATACAACATAAGTGTTGTAAAACGACGGCCAGT | YU1116 |
| TCTTATGTATGTACGTATGTGCTGATTTTTTATGTGCTTGCACAGGAAACAGCTATGACC | YU1117 |
| CCTCTTCCCCACGTCCATTATCAC | YU1347 |
| GTATGTCGATGTTCGATGTTTATGAG | YU1348 |
| GTTCTCTTTTGTTCTCGCTCCCTACATTC | YU1349 |
| ATTGTAGGAGGGCGCGCCAACCTCACG | YU1350 |
| CAAGAACCTTCAAATAACTGGAAATGGGGCATGGAGGATAGCGCCGCAGCTTATCATCGGATCCCCGGGTTAATTAA | YU1518 |
| ATCGGAGAGTATGTATTTGTGTAGTTATGTACTTAGATATGTAACTTAATGAATTCGAGCTCGTTTAAAC | YU1519 |
| TAATGAAGAAAGACATGCAACTAGCAAGAAGAATCAGGGGACAGTTTATTCGGATCCCCGGGTTAATTAA | YU1563 |
| AAACCCCGAAAAAGGGAAAAATCGGCTCCAGCCCTGAAGCACAAATATCAGAATTCGAGCTCGTTTAAAC | YU1564 |
| Plasmid DNA | |
| Plasmid Name | Source |
| pBS-*kanMX4* | Endo Lab |
| pBS-*hphMX* | Endo Lab |
| pFA6a-GFP(S65T)-TRP1 | Longtine et al., 1998 |
| pFA6a-His3MX6-PGAL1 | Longtine et al., 1998 |
| pFA6a-mScarlet-kanMX*4* | Kakimoto et al., 2018 |
| pRS316-su9-RFP | Kakimoto et al., 2018 |
